# A Mechanism of ATF5 Anti-Apoptotic Action that Promotes Leukemia Cells Survival

**DOI:** 10.1101/193292

**Authors:** Stephan P. Persengiev

## Abstract

ATF5 transcription factor was found to play an essential role in hematopoietic cell survival and neuronal cell differentiation. Pro-survival activities of ATF5 are believed to be a result of its ability to inhibit apoptosis. Anti-apoptotic function of ATF5 is almost certainly carried out by regulating gene transcription, but information concerning its target genes has been missing. Here, we report the identification of cyclin D3 as an essential target of ATF5, and provide evidence that its upregulation is critical for the realization of ATF5-induced inhibition of apoptosis. Functional analysis confirmed the association of cyclin D3 with leukemia cells resistance to apoptosis and ATF5-induced splenic dysplasia in transgenic mice. These data link cyclin D3 cell cycle regulatory functions with the apoptotic machinery and reveal a potentially important facet of ATF5 biology.

## Introduction

Mechanisms controlling programmed cell death are deregulated in several human pathologies, most notably cancer and autoimmunity. A critical step in the development of hematopoietic and lymphoid cancers is the abnormal expression pattern of genes that mediate apoptosis. Transcription factors play a critical role in controlling cell proliferation, cell cycle progression and apoptosis (Engel and Murre, 2001; Hickman et al., 2002; Juin et al., 1999). ATF5, a member of ATF/CREB transcription family, has been reported recently to function as an inhibitor of apoptosis in hematopoietic cells and as an essential factor, required for the differentiation of certain neuronal lineages (Angelastro et al., 2005; Mason et al., 2005; Persengiev et al., 2002; Persengiev and Green, 2003). ATF5 expression in neural progenitor cells appears to be necessary for maintaining the proliferation and counteracts the effect of extracellular differentiation factors, such as NGF, NT3 and CNTF (Angelastro et al., 2003; Angelastro et al., 2005). Moreover, ATF5 has been found to be overexpressed in human glioblasmomas and human and rat glioma cell lines (Angelastro et al., 2005). Inhibition of ATF5 activity in these cell resulted in rapid apoptosis suggesting that ATF5 constitutive expression might provide selective survival advantages in these tumors (Sheng et al., 2010; Sheng et al., 2011).

In the immune system, the coordinated expression of stimulatory and inhibitory regulators of programmed cell death is an important ongoing process required for lymphoid development and its disruption is the primary cause for tumor development. Activation of apoptotic/cell death programs usually occurs when cell fail to complete G1/S transition and factors that are differentially expressed during cell cycle progression and interfere with the apoptotic machinery are essential for normal cell development. ATF5 is expressed predominantly during the G1 and G1/S transition points implying that it regulates genes associated with the control of cell cycle progression (Persengiev et al., 2002). Here, we report the identification of cyclin D3 as primary target of ATF5 transcription factor, and provide functional evidence that its upregulation is critical for the realization of ATF5-induced inhibition of apoptosis in hematopoietic cells.

## Results and Discussion

We searched for potential ATF5 transcriptionally regulated genes based on three criteria: a) functional association with cell cycle progression and apoptosis; b) G1 and G1/S restricted expression; and c) presence of ATF/CREB consensus binding sites (TGACGTCA) within the promoter region. E2F1, cyclin A, cyclin D1 and cyclin D3 were identified as probable ATF5 transactivated genes and were subjected to additional in vestigation (Henglein et al., 1994; Herber et al., 1994; Motokura et al., 1992; Shimizu et al., 1998; Slansky et al., 1993; Wang et al., 1996). As a first step in identifying the transcriptional targets of ATF5 we carried out RT-PCR analysis of the selected genes in FL5.12-ATF5 inducible cell derivatives following IL-3 withdrawal. Cyclin D3 was found to be upregulated following ATF5 induction in IL-3-deprived FL5.12-ATF5 cell line (**Fig. 1A)**. In contrast, E2F1, cyclin A and cyclin D1 expression was not affected despite the fact that their promoters contained functional ATF response elements. Moreover, DNA microarray analysis of IL-3 deprived FL5.12-ATF5 cell line identified cyclin D3 as one of the upregulated genes (**Table 1 and Fig. 2**).

**Figure 1.**
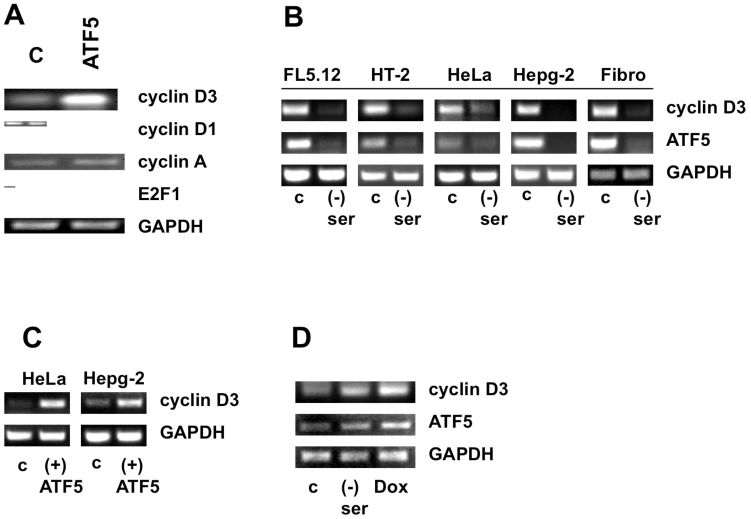
Coordinated expression of cyclin D3 and ATF5 in various cell lines. **a,** RNA isolated from control © and FL5.12-ATF5 (ATF5) cell derivatives were used to analyze the expression of cyclin D3, cyclin D1, cyclin A and E2F1, as well as GAPDH serving as an internal control. **b,** ATF5 and cyclin D3 expression was monitored in various cell lines and primary fibroblasts (fibro) undergoing apoptosis following serum deprivation. **c,** HeLa and Hepg-2 cell line were transiently transfected with an ATF5 expression vector and cyclin D3 mRNA levels were analyzed 48 hours later. **d,** Cyclin D3 and ATF5 expression in HL60/MX2 cells following serum deprivation and doxorubocin treatment. C – control; (-) ser – serum deprived; Dox – doxorubocin treated.

**Table 1.** List of genes with increased or decreased expression in FL5.12-ATF5 cell line after 24 h of serum deprivation.

| Up-regulated genes | Fold increase | Function |
| --- | --- | --- |
| Alix/AIG-2 | 20 | Ca<sup>2+</sup>-binding protein. |
| RAB-14 | 19 | Cell cycle. Proliferation. |
| Cyclin D3 | 16 | Cell cycle. |
| CO-029 antigen | 10 | Carcinogenesis. |
| Jac-1 | 8 | Cell signaling. Proliferation. |
| RAC1 | 8 | Cell cycle. Proliferation. |
| EKLF | 7 | Cell cycle. |
| B-myb | 6 | Proliferation. |
| RAB-1B | 6 | Cell cycle. Proliferation. |
| Lfc | 6 | Ras activator. Oncogene. |
| R-ras | 5 | Cell cycle. Proliferation. |
| Stat2 | 3 | Signal transduction. |
| Stat5b | 3 | Signal transduction. |
| Down-regulated genes | Fold decrease | Function |
| Vigilin | 44 | Lipoprotein-binding factor. |
| Phosphofructokinase | 17 | Glycolysis. |
| FK506 binding protein | 15 | Immunosuppression. |
| Cyclosporin A binding protein | 13 | Immunosuppression. |
| Semaphorin H | 12 | Apoptosis. |
| ATF4 | 10 | Transcription. |
| Synaptophysin | 10 | Apoptotic marker. |
| Cathepsin S | 9 | Lysosomal enzyme. |
| CD24 | 8 | Apoptosis. |
| Calmodulin-like protein NB-1 | 8 | Apoptosis. |

**Figure 2.**
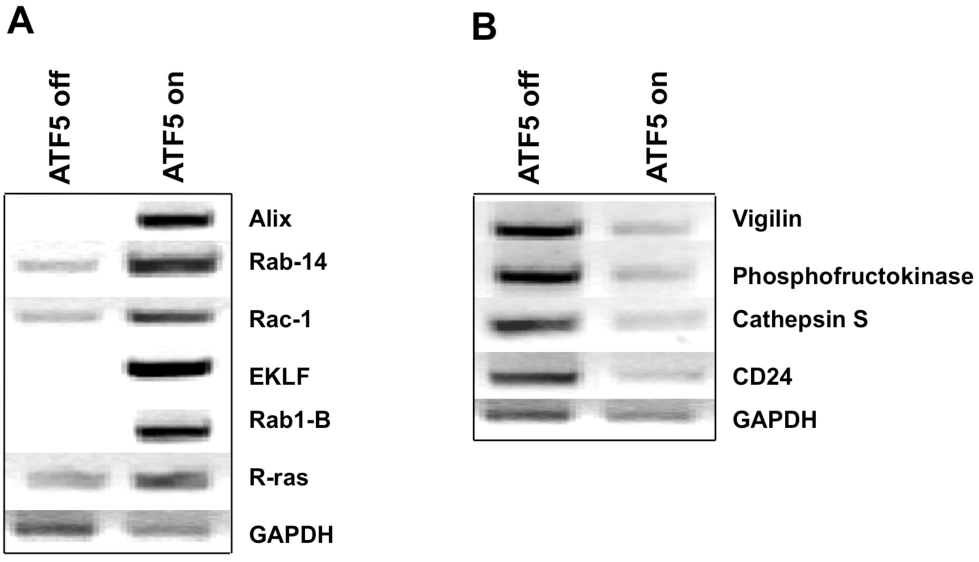
RT-PCR analysis of selected induced and repressed genes in FL5.12-ATF5 cell line. a, upregulated transcripts; b, downregulated transcripts

To confirm that cyclin D3 is a *bona fide* ATF5 regulated gene we performed additional RT-PCRs on various cell lines and tissue samples obtained from transgenic mice expressing ATF5 under the control of lymphocytic-specific Em promoter. Strong correlation was consistently observed only between ATF5 and cyclin D3 expression in several cell lines **(Fig.1B)**. ATF5 and cyclin D3 levels decreased in FL5.12, HT-2, HeLa, Hepg-2 and primary fibroblasts undergoing apoptosis. Analysis of cell lines transiently transfected with an ATF5 expression vector exhibited an increased cyclin D3 expression **(Fig. 1C)**. Cyclin D3 expression levels were relatively low in resting lymphocytes, but significant upregulation was observed in lymphocytes derived from the spleen, lymph nodes and bone marrow of transgenic mice engineered to express ATF5 selectively in hematopoietic cells (**Fig. 3D,** see next paragraph for details). Surprisingly, ATF5 and cyclin D3 expression levels markedly increased in HL-60/MX2 myelocytic leukemia cell line following serum deprivation and doxorubocin treatment **(Fig. 1D)**.

To further understand ATF5 biological functions we generated a hemagglutinin (HA) tagged version of ATF5 protein inserted downstream of lymphocytic-specific EmSR promoter/enhancer construct (Carron et al., 2000). Nine transgenic founder mice for Em-HA-ATF5 construct were obtained. Western blots confirmed that the transgene was selectively expressed in the hematopoietic organs (**Fig. 3A**). Histopathological and cytological analysis revealed that ATF5 founders developed moderate splenomegaly and lymph nodes enlargement. F1 transgenic animals were obtained from two founders (Nos. 23 and 37). Microscopic examination of F1 ATF5 transgenic mice showed disruption in the normal spleen architecture that can be described as a mild hyperplasia, enlargement of the white pulp, and infiltration of lymphoid cells in the red pulp. Lymph nodes exhibited active germinal centers and lymphoid cells infiltration in the peripheral capsule (**Fig. 3B**). The cell surface markers of F1 transgenic animals lymphoid compartment were further characterized by FACS analysis. Lymphocytes from lymph nodes, thymus and spleen and bone marrow were isolated and immunostained with a cohort of lymphocytic-specific PE- or FITC-labeled antibodies. A significant increase of CD4 and CD3/CD28 double-positive lymphocytes was detected in ATF5 transgenic mice (**Table 2**). PCNA proliferation marker staining and annexin V labeling revealed a moderate decrease in the number of lymphoid cells undergoing cell death (**Fig. 3C**).

**Figure 3.**
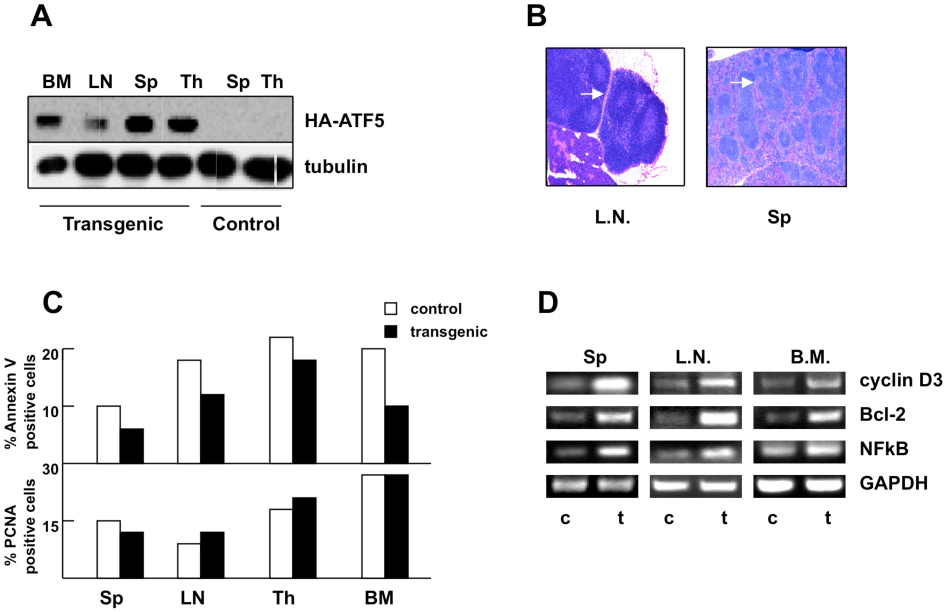
Cyclin D3 expression is upregulated in the hemopoietic organs of ATF5 transgenic mice. **a,** Westen blot analysis of total protein extracts from bone narrow (BM), lymph nodes (LN), spleen (Sp), and thymus (Th) from control and transgenic animals expressing an HA-taged ATF5 transgene under the control of Em lymphocytic-specific enhancer/promoter cassette. Blots were analyzed with sequentially with anti-HA and tubulin antibodies. **b**, Histophology of hematopoietic organs. Lymph nodes are showing lymphocyte infiltration in the peripheral capsule and disruption of normal spleen tissue architecture. Note lymphoid cell infiltration in the spleen red pulp. L.N. – lymph nodes; Sp – spleen. **c,** Flow cytometric detection of Annexin V positive and PCNA expressing lymphoid cells derived from the spleen (Sp), lymph nodes (LN), thymus (Th), and bone marrow (BM). **d,** RT-PCR analysis of cyclin D3, Bcl-2, NFkB and GAPDH transcripts in control © and transgenic (t) mice.

Analysis of potential ATF5 target genes in F1 transgenic mice showed that in addition to the elevated cyclin D3 expression, Bcl-2 and NFkB levels were upregulated in the primary lymphocytes (**Fig. 3D**). These data revealed that ATF5-induced activation of CD28 signaling leads to increased expression of NFkB and Bcl-2 antiapoptotic genes in T-lymphocytes (DeRyckere et al., 2003; Hansen et al., 2002; Tacke et al., 1997; van den Brandt et al., 2004). The mechanism of CD28 activation remains unclear, but most likely is realized by an indirect route. Thus, by interfering with the cyclin D3 and CD28 regulatory cascades, ATF5 alters the lymphoid cell differentiation process and inhibits the apoptotic surveillance mechanisms that limit lymphocytes expansion.

**Table 2.** Cell surface marker analysis of lymphocytic cells obtained from thymus,spleen lymh nodes, and bone marrowof ATF5 transgenic mice. Single-cell suspensions were obtained from the indicated organs of F1 transgenic mice 23-1and 23-2 and analyzed by flow cytometry for expression markers. Age-matched non-transgenic littermates are shown as control.

| **Spleen** | CD4 | CD8 | CD3/CD28 |
| --- | --- | --- | --- |
| Control | 22% | 14% | 35% |
| ATFx-F1 | 26% | 15% | 48% |
| **Thymus** | CD4 | CD8 | CD3/CD28 |
| Control | 9% | 1% | 15% |
| ATFx-F1 | 8% | 1% | 12% |
| **Lymph Nodes** | CD4 | CD8 | CD3/CD28 |
| Control | 36% | 22% | 35% |
| ATFx-F1 | 44% | 26% | 41% |
| **Bone Marrow** | CD4 | CD8 | CD3/CD28 |
| Control | 5% | 2% | 16% |
| ATFx-F1 | 7% | 6% | 20% |

Cyclin D3 upregulation in ATF5 cell derivatives and lymphocytes from transgenic mice suggested that ATF5 might be an activator of cyclin D3 transcription. Previous promoter mapping studies identified putative ATF binding sites in the vicinity of cyclin A, E2F1 and cyclin D1 and D3 promoters (Henglein et al., 1994; Herber et al., 1994; Motokura et al., 1992) (**Fig. 4**). Therefore, to determine whether ATF5 is required for promoter activation to occur we cotransfected an ATF5 expression vector with cyclin A, E2F1, cyclin D1 and cyclin D3 luciferase reporter plasmids in HeLa cells. In parallel, we performed co-transfection experiments with an ATF2 expression construct that has been reported previously to transactivate cyclin A promoter (Shimizu et al., 1998). As demonstrated in **Fig. 5A** transient transfection of ATF5 resulted in a robust activation of cyclin D1 and D3 promoters. In contrast, cyclin A and E2F1 promoters were irresponsive to ATF5. As expected, ATF2 strongly transactivated cyclin A and cyclin D1 expression levels, but had no effect on E2F1 and cyclin D3 transcription. ATF5 specificity was confirmed by employing a dominant-negative ATF5 expression vector (ATF5-DBD), which lacks the transactivation domain (4). As expected, cotransfections of ATF5 and ATF5-DBD led to elimination of ATF5-dependent activation of cyclin D1 and D3 promoters **(Fig. 5A)**.

Based on structural relatedness ATF5 and ATF4 are classified as a separate subgroup of ATF/CREB transcription factors family (Hansen et al., 2002; Vinson et al., 2002). Therefore, we compared the ability of ATF4 and ATF5 to transactivate cyclin D3 promoter. **Figure 5B** shows that ATF4 is even a more potent activator than ATF5 in this transactivation assay. To determine whether ATF5 and ATF4 can synergistically activate cyclin D3 promoter, we cotransfected equimolar amounts of both factors. The results depicted in **Figure 5B** showed a lack of synergism between ATF4 and ATF5. Moreover, it appears that ATF5 is the dominant activator of cyclin D3 transcription based on the comparatively similar luciferase levels in

**Figure 4.**
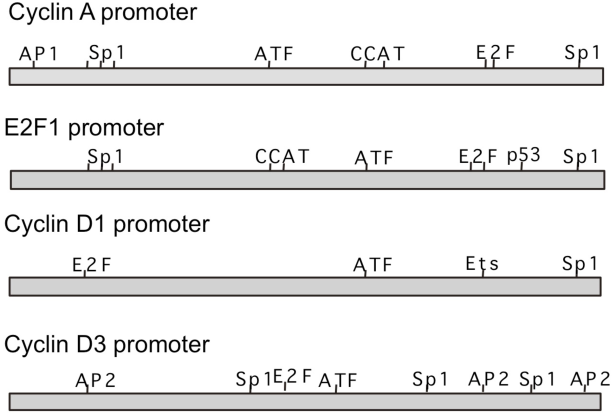
Schematic of cyclin A, E2F1, cyclin D1, and cyclin D3 promoter regions depicting relevant transcription factor binding sites

**Figure 5.**
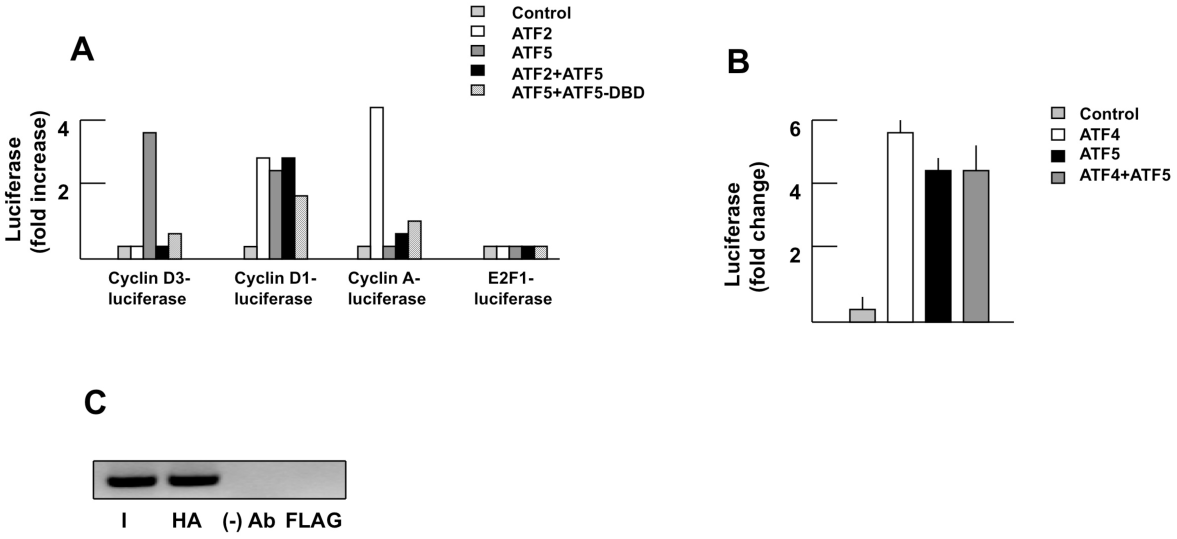
ATF5 specifically activates cyclin D3 promoter. **a,** Promoter analysis. HeLa cells were cotransfected with cyclin D3, cyclin D1, cyclin A, and E2F1 reporter constructs and either an empty vector or plasmids directing the expression of ATF2, ATF5, and ATF5-DBD. Normalized promoter activity was determined 48 h later. Results are means of at least two independent experiments each with triplicate determinations. **b,** Cyclin D3 reporter constuct was cotransfected with ATF4 and ATF5 expression vectors and luciferase activity was determined 48 h later. **c,** Chromatin immunoprecipitation (ChIP) analysis of ATF5 association with cyclin D3 promoter in vivo in FL5.12-ATF5 cells (upper panel) and mouse splenocytes isolated from transgenic mice (lower panel). Anti-HA antiserum was used to precipitate the chromatin-bound ATF5, as well as a FLAG antibody was employed a negative control.

ATF5 alone and ATF4/ATF5 combined transfections (**Fig. 5B**). Finally, we confirmed ATF5 association with cyclin D3 promoter elements *in vivo* in FL5.12-ATF5 cell line and slenocytes from ATF5 transgenic mice by chromatin immunoprecipitation (ChIP) employing an HA-tag antibody (**Fig. 5C**).

The persistent cyclin D3 expression in apoptotic-resistant HL-60 cells implied that cyclin D3 promotes cell survival. To address this question, we tested the effect of ectopically expressed cyclin D3 on the rate of apoptosis in growth factor deprived FL5.12 and HL-60 cell lines. We derived FL5.12 and HL-60 cell lines stably expressing cyclin D3 cDNA under the control of an inducible promoter. **Figure 6A** shows that the expression of cyclin D3 was induced in the absence of doxycycline in HL-60 and FL5.12 cell derivatives. As expected FL5.12 cells underwent rapid apoptosis following IL-3 withdrawal, whereas FL5.12-cyclin D3 derivatives displayed significant resistance to apoptosis (**Fig. 6B**). HL-60 cells were resistant to apoptosis when deprived of serum, and exogenous cyclin D3 expression had no detectable effects on cell death and cell proliferation rate (Boland et al., 2000; Jones et al., 2002) (**Fig. 6C**). To ask whether cyclin D3 expression might have a more subtle effect on cell growth, we used 3H-thymidine labeling and DNA flow cytometry to compare more accurately the proliferation rate and cell cycle progression in asynchronous populations of HL-60-cyclin D3 cell line. The number of actively dividing cells and G1, S and G2/M phases were comparable in the presence or absence of overexpressed cyclin D3, indicating that cell cycle progression was not altered (**Fig. 6D and E**).

**Figure 6.**
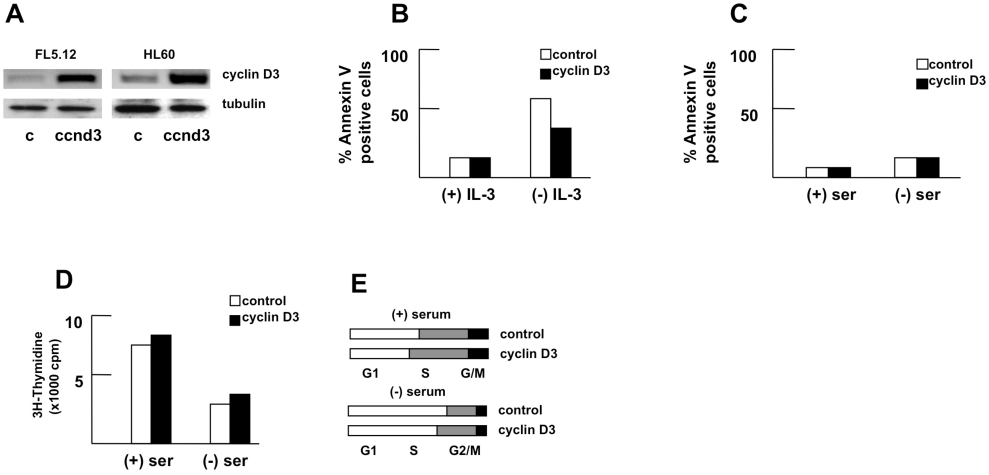
Cyclin D3 induction in HL-60 cells does not promote cell proliferation. **a,** Expression analysis of cyclin D3 in FL5.12 and HL-60 cells stably expressing cyclin D3. Whole-cell extracts prepared from control and cyclin D3 cell derivatives (ccnd3) were probed sequentially with cyclin D3 and tubulin antiserums. **b,** Cyclin D3 overexpression in FL5.12 cell inhibits apoptosis. Wild-type and cyclinD3-FL5.12 cells were cultured in the presence and absence of IL-3 and percentage of apoptotic cells was monitored by annexin V staining. **c,** Annexin V staining of HL-60-cyclin D3 cell derivatives following serum deprivation. **d,** DNA synthesis rate analysis of HL60-cyclin D3 cells by ^3^H-thymidine incorporation assay. **e,** Cell cycle profiling of HL-60-cyclin D3 cells cultured in the presence and absence of serum. The percentage of cells in each cell cycle phase in an asynchronous cell population was determined by FACS analysis. The relative numbers of cells in G^0^/G^1^, S, and G^2^/M phases are shown schematically.

Next, we tested whether cyclin D3 is required for cell survival and resistance to apoptosis. **Figure 7** depicts the results of knockdown experiments using siRNAs directed against the cyclin D3 mRNA. HL-60 cells cultured in the presence or absence of serum were transiently transfected with either cyclin D3 or non-specific siRNAs for 48 hours. Cyclin D3 levels were reduced by approximately 90% following cyclin D3 siRNA application (**Fig 7A)**. As expected, non-specific siRNA has no effect on cyclin D3 expression. Annexin V staining on HL-60 cells revealed that cyclin D3 inhibition increased the susceptibility of HL-60 cells to apoptosis following serum deprivation (**Fig. 7B**). However, resistance to treatment with the chemotherapeutic drug doxorubicin was not affected (data not shown).

Also, we tested whether the increased expression of cyclin D3 in serum deprived HL-60 cells leads to alteration in the expression pattern of cdk4, cdk6 and pRb proteins. Western blotting showed that pRb levels decreased substantially following serum withdrawal, while cdk4 and cdk6 remained stable. Cyclin D3 siRNA treatment did not cause any detectable changes in pRb and cdk4 and cdk6 expression under these experimental conditions (**Fig. 8**). Thus, the anti-apoptotic activities of cyclin D3 were not associated with pRb/E2F1 regulated transcription cascade.

The mechanisms by which constitutive ATF5 expression leads to inhibition of apoptosis and as a consequence to uncontrolled proliferation in hematopoietic cells remain to be fully explored. Our previous findings suggested that ATF5 is able to block the “extrinsic” apoptotic pathway by interfering with the signal cascades initiated by death receptor activation by acting on downstream targets (Persengiev et al., 2002). Interestingly, ATF5 has been reported to be in a physical complex with cyclin D3 (Liu et al., 2004). Cyclin D3 association with ATF5 appears to augment ATF5 transcription activity. Thus, in line with the studies presented here, it is likely that the constitutive ATF5 expression leads to cyclin D3 accumulation, which in turn is capable of maintaining its own expression by stabilizing ATF5 transcription activity. These findings raise the possibility that ATF5 and cyclin D3 may form a functional self-sustained positive feedback loop that operates in certain tumor cells and facilitates their survival.

**Figure 7.**
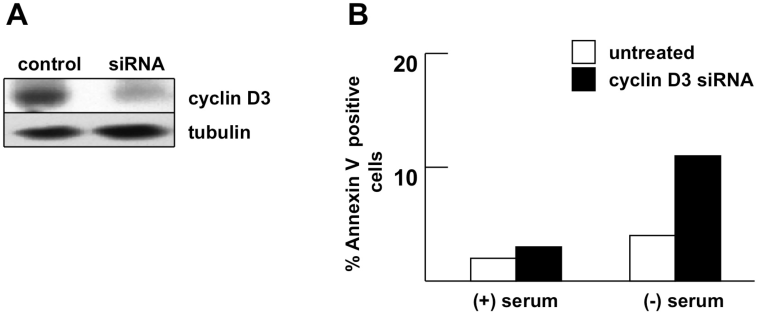
Cyclin D3 expression is required for cell survival in HL-60 cells **a,** Analysis of siRNA-induced cyclin D3 down-regulation in HL-60 cells. Whole-cell protein extracts prepared from control and siRNA treated cells were probed with cyclin D3 and tubulin antibodies. **b,** Annexin V staining of HL-60 cells treated with cyclin D3 siRNA. Percentage of apoptotic cells was evaluated after 48 h of serum deprivation.

**Figure 8.**
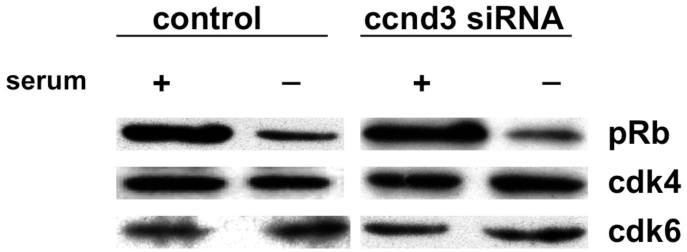
Western blot analysis of pRB, cdk4, and cdk 6 in HL60 cells following serum deprivation.

In conclusion, we identified cyclin D3 as an important target of ATF5 transcription factor and provided functional evidence for its association with leukemia cell resistance to apoptosis. On the basis of these results, we propose the following model for the biological role of ATF5 in hematopoietic cells and its relevance for lymphoproliferative disease development. Under normal conditions ATF5 serves as the principal activator of cyclin D3 transcription and is restricted to G1 and G1/S cell cycle phases. Constitutive and/or over-expression of ATF5 lead to alteration in the expression signature of cyclin D3, which is no longer limited to G1 phase. The biological consequence of this cascade is the acquired resistance to apoptotic signals. An implication of this model is that altered cyclin D3 expression and deactivation of apoptotic surveillance mechanisms might serve as an oncogenic event in hematopoietic cells

## Materials and Methods

### Cells culture and transfection

HL-60/MX and FL5.12 cells were propagated at 37 C in DMEM (Life Technologies) supplemented with 10% fetal bovine serum, 100 U/ml penicillin and 100 U/ml streptomycin. Cells were regularly passed to maintain exponential growth. Transfections with different expression vectors were carried out by using Fugene 6 (Roche Applied Science) according to manufacturer’s instructions. Cyclin D3 siRNA SMARTpool (Dharmacon) transfections were performed by Oligofectamine (Life Technologies) and 100 nM siRNA mixture, as recommended by the manufacturer.

### Microarray analysis

Total RNA was prepared from control FL5.12 cell line and FL5.12-ATF5 cell derivative following 24h of serum deprivation by RNeasy kit (Qiagen). Targets for hybridization to the microarrays were prepared as described (Devireddy et al., 2005). Hybridization and scanning of mouse Mu11K GeneChip arrays (Affymetrix) were performed as recommended by the manufacturer.

### Transgenic mice

ATF5 expression vector was constructed by subcloning the full-length mouse ATF5 cDNA containing a hemagglutinin (HA) tag at the N terminus into EmSRa vector (Bodrug et al., 1994; Carron et al., 2000). This vector drives the expression of exogenous cDNA from the EmSRa enhancer/promoter cassette selectively in hematopoietic cells. The EmSRa-HA-ATF5 construct was digested by NotI to release the transgene before injection into fertilized eggs.

Transgenic C57B6 animals were generated by pronuclear microinjections at the University of Massachusetts Transgenic Core Facility as described previously (Carron et al., 2000). Transgenic mice were identified by polymerase chain reaction (PCR) analysis of tail DNA using primers spanning the HA-tag and ATF5 cDNA transgene regions.

### RNA and protein analysis

RT-PCR analysis was performed according to standard protocol with total RNA prepared from cultured cell lines and tissue samples by Trizol (Roche Applied Science).

**Table 3.**
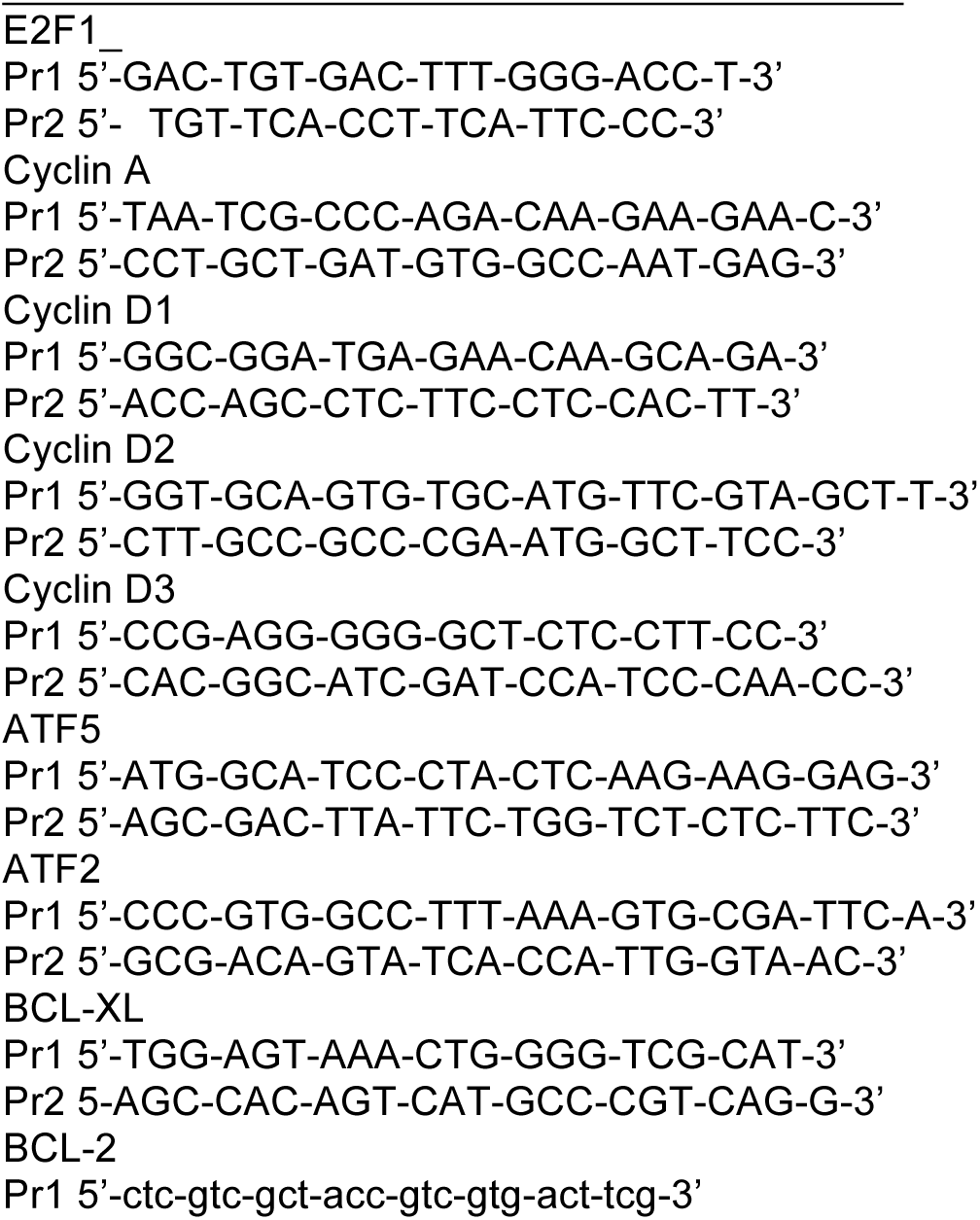
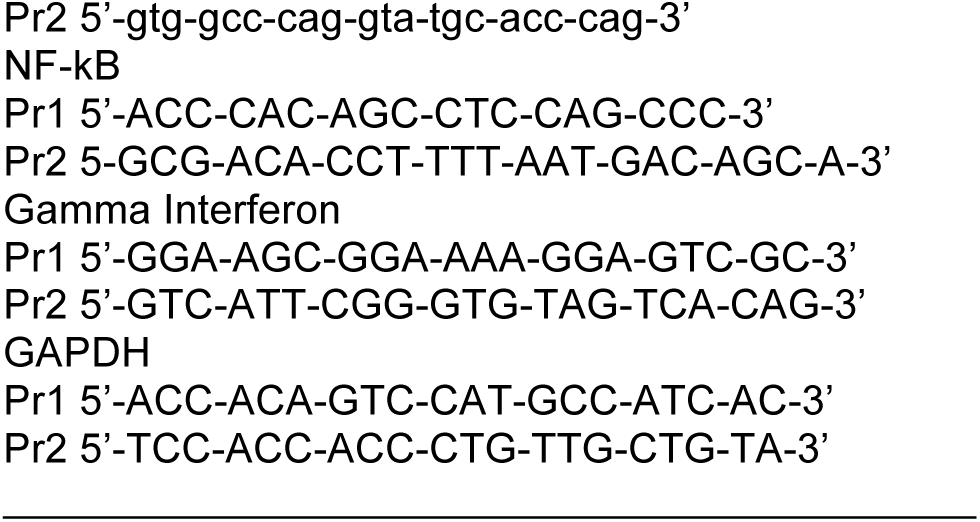
Primer sequences used for RT-PCR analysis are listed below.

Immunoblot assays were performed using polyclonal antibodies that specifically recognized the HA tag, cyclin D3, pRb, cdk 4, cdk 6 (Santa Cruz Biotechnology) and ATF5 (Novus Biologicals).

### Histopathological and flow cytometric analysis

Mice were fully dissected with microscopic analysis of all organs. Spleens, livers and thymuses were weighed. All organs were fixed in 10% formaldehyde for 2 hours and further processed for paraffin embedding. Sections were stained with hematoxylin-eosin (HE) and histologically analyzed. Single-cell suspensions from lymph nodes, spleen, thymus, and bone marrow were prepared in RPMI medium containing 10% FCS. Aliquotes of 10^6^ cells were resuspended in PBS containing 3% FCS and processed for immunostaining as previously described (Castilla et al., 1999; Schwaller et al., 2000). Monoclonal antibodies specific for CD4, CD8, CD3, CD28, CD90, CD95, CD45/B220, CD11, and PCNA conjugated with either fluorescein isothiocyanate (FITC) or phycoerythrin (PE) (BD Biosciences) were used. Cytometric analysis was performed on FACsCalibur (Becton Dickinson). A total of 20 000 events were acquired using CellQuest software (Becton Dickinson).

### Chromatin immunoprecipitation (ChIP) assays

The amount of HA-ATF5 bound to the endogenous cyclin D3 promoter in FL5.12-HA-ATF5 cells and splenocytes isolated from ATF5 transgenic mice was measured by ChIP assay, using a HA tag antibody as previously described (Weinmann and Farnham, 2002). Primer sequences used for the PCR amplification of cyclin D3 promoter fragments containing ATF/CREB binding site are: Pr1; 5’-GAG-CCC-AGC-GGA-CCA-ATG-G-3’ and Pr2; 5’-GTA-GCA-ACC-GTG-GAA-TGC-TC-3’

### Apoptotic and cell cycle analysis

Apoptosis was analyzed by staining with annexin V (Oncogene Research Products) according to manufacturer instructions. Cell cycle analysis was performed on asynchronously growing cells. Cellular DNA was stained with propidium iodide and DNA content and cell number analyzed by FACS analysis. Cell cycle profiles were analyzed using ModFit program (Verify Software).

## Acknowledgments

The experimental results reported in this manuscript were designed by Dr. Stephan Persengiev and performed in the Program in Gene Function & Expression, University of Massachusetts Medical School, Worcester, MA, USA The author is grateful to Michael Green and members of his lab for help, insights and suggestions throughout the work. This work was supported by an American Cancer Society grant to SPP.

## References

Angelastro, J.M., Ignatova, T.N., Kukekov, V.G., Steindler, D.A., Stengren, G.B., Mendelsohn, C., and Greene, L.A. (2003). Regulated expression of ATF5 is required for the progression of neural progenitor cells to neurons. J Neurosci 23, 4590–4600.

Angelastro, J.M., Mason, J.L., Ignatova, T.N., Kukekov, V.G., Stengren, G.B., Goldman, J.E., and Greene, L.A. (2005). Downregulation of activating transcription factor 5 is required for differentiation of neural progenitor cells into astrocytes. J Neurosci 25, 3889–3899.

Bodrug, S.E., Warner, B.J., Bath, M.L., Lindeman, G.J., Harris, A.W., and Adams, J.M. (1994). Cyclin D1 transgene impedes lymphocyte maturation and collaborates in lymphomagenesis with the myc gene. EMBO J 13, 2124–2130.

Boland, M.P., Fitzgerald, K.A., and O'Neill, L.A. (2000). Topoisomerase II is required for mitoxantrone to signal nuclear factor kappa B activation in HL60 cells. J Biol Chem 275, 25231–25238.

Carron, C., Cormier, F., Janin, A., Lacronique, V., Giovannini, M., Daniel, M.T., Bernard, O., and Ghysdael, J. (2000). TEL-JAK2 transgenic mice develop T-cell leukemia. Blood 95, 3891–3899.

Castilla, L.H., Garrett, L., Adya, N., Orlic, D., Dutra, A., Anderson, S., Owens, J., Eckhaus, M., Bodine, D., and Liu, P.P. (1999). The fusion gene Cbfb-MYH11 blocks myeloid differentiation and predisposes mice to acute myelomonocytic leukaemia. Nat Genet 23, 144–146.

DeRyckere, D., Mann, D.L., and DeGregori, J. (2003). Characterization of transcriptional regulation during negative selection in vivo. J Immunol 171, 802–811.

Devireddy, L.R., Gazin, C., Zhu, X., and Green, M.R. (2005). A cell-surface receptor for lipocalin 24p3 selectively mediates apoptosis and iron uptake. Cell 123, 1293–1305.

Engel, I., and Murre, C. (2001). The function of E- and Id proteins in lymphocyte development. Nat Rev Immunol 1, 193–199.

Hansen, M.B., Mitchelmore, C., Kjaerulff, K.M., Rasmussen, T.E., Pedersen, K.M., and Jensen, N.A. (2002). Mouse Atf5: molecular cloning of two novel mRNAs, genomic organization, and odorant sensory neuron localization. Genomics 80, 344–350.

Henglein, B., Chenivesse, X., Wang, J., Eick, D., and Brechot, C. (1994). Structure and cell cycle-regulated transcription of the human cyclin A gene. Proc Natl Acad Sci U S A 91, 5490–5494.

Herber, B., Truss, M., Beato, M., and Muller, R. (1994). Inducible regulatory elements in the human cyclin D1 promoter. Oncogene 9, 2105–2107.

Hickman, E.S., Moroni, M.C., and Helin, K. (2002). The role of p53 and pRB in apoptosis and cancer. Curr Opin Genet Dev 12, 60–66.

Jones, K.H., Liu, J.J., Roehm, J.S., Eckel, J.J., Eckel, T.T., Stickrath, C.R., Triola, C.A., Jiang, Z., Bartoli, G.M., and Cornwell, D.G. (2002). Gamma-tocopheryl quinone stimulates apoptosis in drug-sensitive and multidrug-resistant cancer cells. Lipids 37, 173–184.

Juin, P., Hueber, A.O., Littlewood, T., and Evan, G. (1999). c-Myc-induced sensitization to apoptosis is mediated through cytochrome c release. Genes Dev 13, 1367–1381.

Liu, W., Sun, M., Jiang, J., Shen, X., Sun, Q., Liu, W., Shen, H., and Gu, J. (2004). Cyclin D3 interacts with human activating transcription factor 5 and potentiates its transcription activity. Biochem Biophys Res Commun 321, 954–960.

Mason, J.L., Angelastro, J.M., Ignatova, T.N., Kukekov, V.G., Lin, G., Greene, L.A., and Goldman, J.E. (2005). ATF5 regulates the proliferation and differentiation of oligodendrocytes. Mol Cell Neurosci 29, 372–380.

Motokura, T., Keyomarsi, K., Kronenberg, H.M., and Arnold, A. (1992). Cloning and characterization of human cyclin D3, a cDNA closely related in sequence to the PRAD1/cyclin D1 proto-oncogene. J Biol Chem 267, 20412–20415.

Persengiev, S.P., Devireddy, L.R., and Green, M.R. (2002). Inhibition of apoptosis by ATFx: a novel role for a member of the ATF/CREB family of mammalian bZIP transcription factors. Genes Dev 16, 1806–1814.

Persengiev, S.P., and Green, M.R. (2003). The role of ATF/CREB family members in cell growth, survival and apoptosis. Apoptosis 8, 225–228.

Schwaller, J., Parganas, E., Wang, D., Cain, D., Aster, J.C., Williams, I.R., Lee, C.K., Gerthner, R., Kitamura, T., Frantsve, J., et al. (2000). Stat5 is essential for the myelo- and lymphoproliferative disease induced by TEL/JAK2. Mol Cell 6, 693–704.

Sheng, Z., Li, L., Zhu, L.J., Smith, T.W., Demers, A., Ross, A.H., Moser, R.P., and Green, M.R. (2010). A genome-wide RNA interference screen reveals an essential CREB3L2-ATF5-MCL1 survival pathway in malignant glioma with therapeutic implications. Nat Med 16, 671–677.

Sheng, Z., Ma, L., Sun, J.E., Zhu, L.J., and Green, M.R. (2011). BCR-ABL suppresses autophagy through ATF5-mediated regulation of mTOR transcription. Blood 118, 2840–2848.

Shimizu, M., Nomura, Y., Suzuki, H., Ichikawa, E., Takeuchi, A., Suzuki, M., Nakamura, T., Nakajima, T., and Oda, K. (1998). Activation of the rat cyclin A promoter by ATF2 and Jun family members and its suppression by ATF4. Exp Cell Res 239, 93–103.

Slansky, J.E., Li, Y., Kaelin, W.G., and Farnham, P.J. (1993). A protein synthesis-dependent increase in E2F1 mRNA correlates with growth regulation of the dihydrofolate reductase promoter. Mol Cell Biol 13, 1610–1618.

Tacke, M., Hanke, G., Hanke, T., and Hunig, T. (1997). CD28-mediated induction of proliferation in resting T cells in vitro and in vivo without engagement of the T cell receptor: evidence for functionally distinct forms of CD28. Eur J Immunol 27, 239–247.

van den Brandt, J., Wang, D., and Reichardt, H.M. (2004). Resistance of single-positive thymocytes to glucocorticoid-induced apoptosis is mediated by CD28 signaling. Mol Endocrinol 18, 687–695.

Vinson, C., Myakishev, M., Acharya, A., Mir, A.A., Moll, J.R., and Bonovich, M. (2002). Classification of human B-ZIP proteins based on dimerization properties. Mol Cell Biol 22, 6321–6335.

Wang, Z., Sicinski, P., Weinberg, R.A., Zhang, Y., and Ravid, K. (1996). Characterization of the mouse cyclin D3 gene: exon/intron organization and promoter activity. Genomics 35, 156–163.

Weinmann, A.S., and Farnham, P.J. (2002). Identification of unknown target genes of human transcription factors using chromatin immunoprecipitation. Methods 26, 37–47.

